# African Swine Fever Virus Protein B263R Inhibits Autophagy by Promoting Proteasomal Degradation of Beclin1

**DOI:** 10.1101/2024.04.09.588710

**Authors:** Shuxiang Ding, Xingbo Wang, Tingjuan Deng, Kang Xu, Weiren Dong, Yan Yan, Jiyong Zhou, Boli Hu

## Abstract

African swine fever (ASF) is a highly contagious viral disease that affects both domestic and wild pigs, leading to substantial economic losses in the swine industry. However,the precise role of ASFV-encoded viral protein in viral replication and disease has remained elusive. In our study, we elucidated that B263R inhibited autophagy by interacting with Beclin1. Mechanistically, the binding of B263R to Beclin1 promotes its degradation through the mediation of K48-linked ubiquitination at site K31. Notably, the knockdown of Beclin1 mitigated the effect of B263R on reducing the levels of LC3-II, thus further affirming the pivotal role of Beclin1 in B263R-mediated autophagy inhibition. The proliferation of recombinant adenovirus inherently expressing B263R significantly decreased, underscoring the inherent role of B263R-mediated autophagy in virus replication. These findings shed light on the molecular mechanisms employed by ASFV to evade host cell autophagic defenses and offered potential targets for therapeutic interventions against ASFV infection.

**IMPORTANCE:** Understanding the mechanisms underlying the evasion of host cellular defenses by the African swine fever virus (ASFV) is of pivotal significance owing to its profound impact on the global swine industry. The precise functional roles of ASFV proteins in virus replication and disease remain incompletely characterized. In this study, it is demonstrated that a specific ASFV protein, B263R, acts to inhibit autophagy by interacting with Beclin1, thereby facilitating its degradation. The elucidation of this molecular pathway not only advances our comprehension of ASFV pathogenesis but also furnishes potential targets for therapeutic interventions against ASFV infection. By unraveling the intricate interplay between ASFV and host cell autophagy, this scholarly endeavor contributes invaluable insights to the development of strategies aimed at mitigating the spread of ASFV and minimizing its adverse repercussions on the swine industry.

## Introduction

African swine fever (ASF) is a devastating viral disease caused by the infectious African swine fever virus (ASFV), leading to high morbidity and mortality in affected pig populations. This disease has inflicted significant damage on the global pork industry (1). ASFV belongs to the sole member of the family Asfarviridae and primarily replicates in the cytoplasm of macrophages (2). It is characterized by a double-stranded (ds) linear DNA genome enclosed in an enveloped nucleoplasmic exterior, containing more than 151 open reading frames (ORFs) (3). Despite its longstanding impact, the development of a reliable vaccine against ASFV has remained elusive for over a century. This challenge stems from a lack of comprehensive research into ASFV pathogenesis and a limited understanding of the innate immune evasion strategies employed by the virus (4). Efforts to overcome these obstacles are crucial for the development of secure and effective vaccines to combat ASF and mitigate its detrimental effects on the swine industry.

Autophagy is involved in the degradation of organelle and protein during the process of cancer, microbial infection, neurodegeneration, ageing, etc.(5). Many proteins, such as Beclin1(BECN1), collaboratively regulate the autophagy process(6). BECN1 serves as a critical component of Class III phosphoinositide 3-Kinase (PI3K) complexes, which generate phosphorylated phosphatidylinositol (PtdIns) to facilitate the maturation of autophagosomes (7). Among the various post-translational modifications of BECN1, ubiquitination stands out as one of the most significant (8, 9). Recent studies have highlighted the role of BECN1 ubiquitination in the viral life cycle. For instance, during the proliferation of the Middle East Respiratory Syndrome Coronavirus, there is a reduction in the level of ubiquitinated BECN1, leading to the inhibition of autophagosome-lysosome fusion (10). Additionally, USP19 modulates the ubiquitination process of BECN1, promoting autophagosome formation while concurrently suppressing DDX58/RIG-I-mediated type I interferon signaling (11).

Autophagy plays a central role in host defense against viral infection (12). It has been documented that ASFV possesses the capability to manipulate autophagy and hinder autophagosome formation through distinct mechanisms (13). Specifically, ASFV A179L disrupts autophagosome formation by interacting with BECN1(14). On the other hand, K205R activates autophagy via the AKT/mTOR/ULK1 signaling pathway (15). Moreover, E199L also activates complete autophagy by down-regulating the expression of PYCR2 (16). Additionally, A137R promotes the autophagy-mediated lysosomal degradation of TBK1, leading to a reduction in IFN production (17). The process of p17-mediated mitophagy led to the breakdown of mitochondrial proteins involved in antiviral signaling and suppressed the synthesis of IFN-α, IL-6, and TNFα (18).

Previous research showed that B263R has the characteristics of a TATA binding protein and is therefore likely to be involved in viral gene transcription (19). However, effect of B263R in life cycle of ASFV is not investigated in detail. In our current investigation, we discovered that ASFV protein B263R hinders autophagy and facilitates the degradation of BECN1 via the ubiquitination-proteasomal pathway. Moreover, we found that K48-linked ubiquitination of BECN1 at the K31 sites is crucial for B263R-mediated ubiquitination. The result of adenovirus replication showed that B263R substantially suppresses virus proliferation. This study unveils a novel function of B263R-mediated autophagy in immune evasion by ASFV.

## Results

### ASFV B263R inhibits autophagy

To identify the key protein of ASFV on regulating autophagy, we tested the autophagy by respectively transfecting eleven plasmids of ASFV-encoding protien and an empty vector into HEK-293T cells and detecting the levels of SQSTM1/p62 (an autophagy receptor for degradation of substrates). The results showed that B263R leads to an increase in p62 (Figure S1). We further investigated the regulatory function of B263R in autophagy by detecting SQSTM1 and LC3 in PK-15 cells. As shown in Figure 1A-B, the level of SQSTM1 increased greatly, while the conversion of LC3-I to LC3-II decreased significantly, indicating that B263R inhibited autophagy. We then determined the effect of B263R on autophagy stimulation. the similar results revealed in culture environment of Earle’s Balanced Salt Solution (EBSS). Additionally, we also investigated whether B263R prohibited the formation of GFP-LC3 puncta to signify the development of autophagosomes. Figure 1C-D showed that B263R expression significantly reduced the number of GFP-LC3 puncta in 3D4/21 cells with rapamycin treatment. The above data suggested that B263R inhibited autophagy.

**Figure 1.**
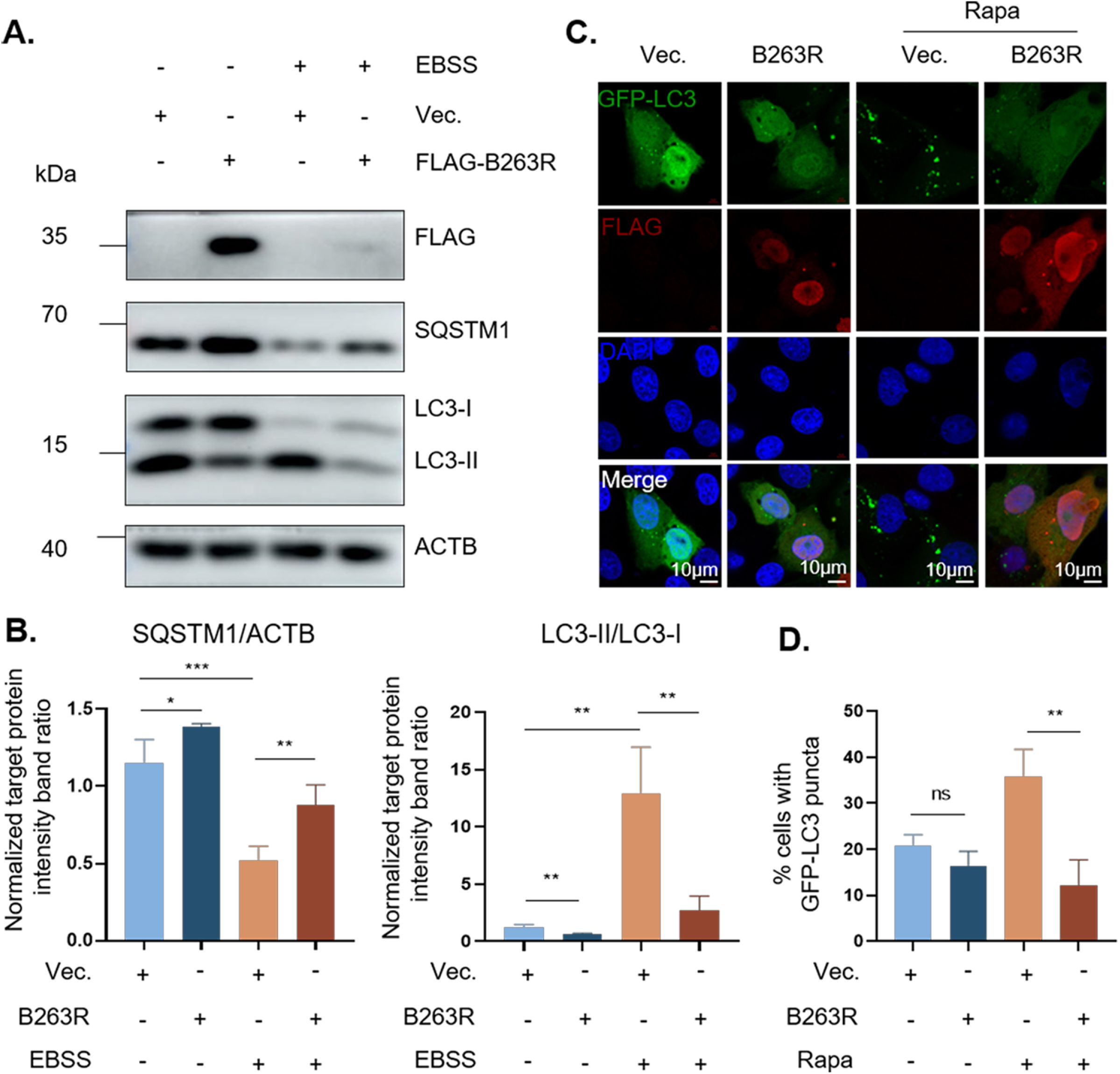
ASFV B263R inhibits autophagy. (A) PK-15 cells were transfected with empty vectors or plasmids expressing Flag-tagged ASFV B263R. Cells were treated with starvation medium (EBSS) for 6h. At 36h post-transfection, cells were harvested for WB analysis by using anti-Flag, anti-SQSTM1, anti-LC3, and anti-ACTB antibodies. (B) Densitometric analysis of protein level by using Image J software, the gray value of ACTB as the control standard. (C) 3D4/21 cells were co-transfected with vectors expressing FLAG or FLAG-B263R and GFP-LC3 for 24h. Cells were treated with Rapamycin (5nM) for 4h. The resultant cells are subjected to confocal microscopy and immunofluorescence staining Scale bar: 5 μm. (D) The numbers of puncta were counted. Error bars: Mean ± SD of 3 independent tests. two-tailed Student’s test; *p < 0.05; **p < 0.01; ***p < 0.001 compared to control.

### The proteasomal degradation of BECN1 is medicated by the interaction with B263R

To examine the mechanisms underlying the pB263R-mediated inhibition of autophagy, we assessed whether the B263R-induced inhibition occurs during autophagosome formation by detecting the change of Phosphatidylinositol 3-phosphate (PI3P), which is crucial for the initial biogenesis of autophagosome (20). Intracellular PI3P contains FYVE domain (21), we therefore examined the distribution of intracellular GFP-2×FYVE puncta. As shown in Figure 2A, there was a obvious reduction of 2×FYVE puncta in B263R overexpressed cells. In order to seek the potential host proteins that interacted with B263R, co-immunoprecipitation (Co-IP) assay was performed to evaluate the interaction between B263R and components of Class III PI3K complex, including the core proteins Vps34, UVRAG, ATG14 and BECN1. Our results indicated that B263R interacted with Belcin1, but not with Vps34, UVRAG, or ATG14 (Figures 2B and S2). Moreover, confocal microscopy revealed a colocalization between B263R and BECN1, when 3D4/21 cells were co-transfected with GFP-B263R and Flag-BECN1 plasmids, Pearson’s Coefficient: 0.799 (Figure 2C), indicating that B263R interacted with the host protein BECN1. We then tested effect of B263R on BECN1, Flag-B263R was transfected into 293T cells in an increasing dose and the results showed that the level of BECN1 displayed a downward trend (Figure 2D). As shown in Figure 2E, the declining BECN1 recovered when treated with proteasome inhibitor MG132. However, BECN1 did not cause a further increase when autophagy was blocked by CQ (Figure 2F). The above results indicated that B263R mediated the proteasomal degradation of Beclin1.

**Figure 2.**
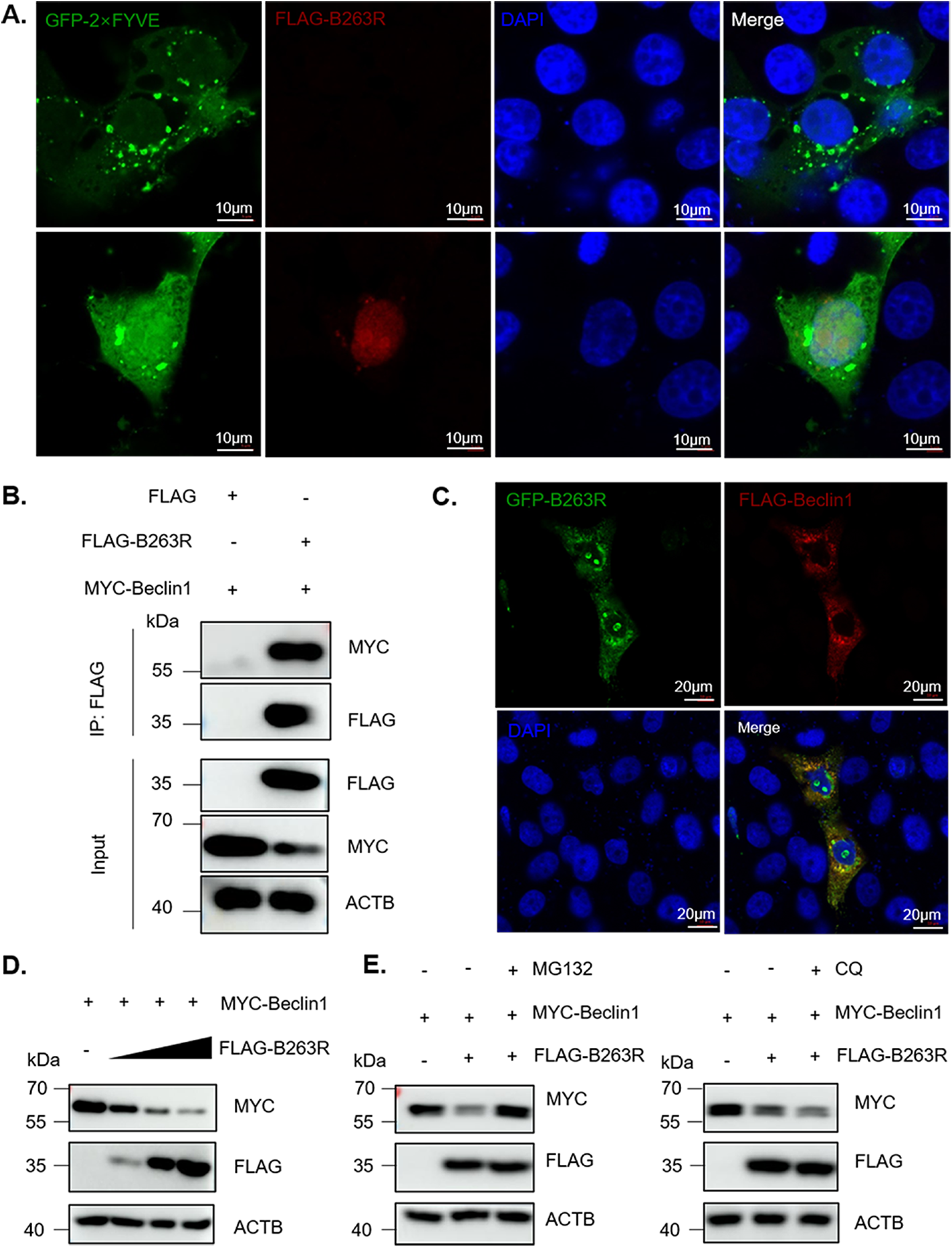
The proteasomal degradation of BECN1 is medicated by the interaction with B263R. (A) 3D4/21 cells were co-transfected with vectors expressing GFP-2×FYVE and FLAG-B263R for 24h. The resultant cells are subjected to confocal microscopy and immunofluorescence staining, scale bars: 10 µm. (B) HEK-293T cells were co-transfected with vectors expressing FLAG or FLAG-B263R and MYC-BECN1 for 36h. At 36h post-transfection, cells were harvested for Co-IP. Immunobloting analysis was performed with anti-FLAG, anti-MYC, and anti-ACTB antibodies. (C) 3D4/21 cells were co-transfected with vectors expressing GFP-B263R and FLAG-BECN1 for 24h. The fluorescence signals were visualized by confocal immunofluorescence microscopy, scale bars: 20 µm. (D) B263R promotes degradation of BECN1 in a dose-dependent manner. Increasing amounts of B263R-expressing plasmids and MYC-BECN1 were co-transfected into HEK-293T cells, and the BECN1 was analyzed by detecting the protein levels. (E) HEK293T cells were co-transfected with vectors expressing FLAG or FLAG-B263R and MYC-BECN1 for 30 h, followed by treatment with MG132 (20μM) or chloroquine phosphate (50μM) for 6h.

### The BECN1 K31 residue is essential for B263R-mediated K48-ubiquitination

Ubiquitin (Ub) binds to the lysine (K) residue of substrate proteins through an enzyme cascade, including E1 activators, E2 binders, and E3 ubiquitin ligases (22). To determine whether B263R facilitated the ubiquitination-dependent degradation of BECN1, we assessed the level of poly-ubiquitin linkages on BECN1 in the presence or absence of B263R. As illustrated in Figure 3A, B263R overexpression led to significant enhancement in BECN1 ubiquitination. To investigate the major site of B263R-mediated ubiquitination, we constructed three functional domains plasmids of BECN1 and confirmed the interacting domain by Co-IP. The result, illustrated in Figure 3B, suggested that B263R actually interacted with N-terminal of BECN1. Furthermore, the cells overexpressing the functional domain is co-transfected with Flag-B263R in an increasing dose and observed the level of N-terminal, coiled-coil (CC) domain, β-α autophagy-specific (BARA) domain. B263R indeed promoted degradation of N-terminal segment of BECN1 in a dose-dependent manner (Figure 3C). However, the promotion of B263R on ubiquitination of BECN1 N-terminal with five lysines was nearly entirely abrogated upon substitution of K31 with arginine (Figure 3D-3E). Moreover, it was evident that the reduction observed in the B263R K^31^R mutant was not as pronounced in the presence of the B263R K^25^R mutant (Figure 3F). Given that the specific ubiquitin chain attached to the lysine residues of the substrate protein triggers unique biological responses(23), we detected ubiquitin linkage types (K6, K11, K27, K29, K33, K48, K63) conjugated onto BECN1 driven by B263R. The results showed that K48 ubiquitin may be linked to BECN1(Figure 3G). Overall, these results indicated that the BECN1 K31 residue was essential for B263R-mediated K48-ubiquitination.

**Figure 3.**
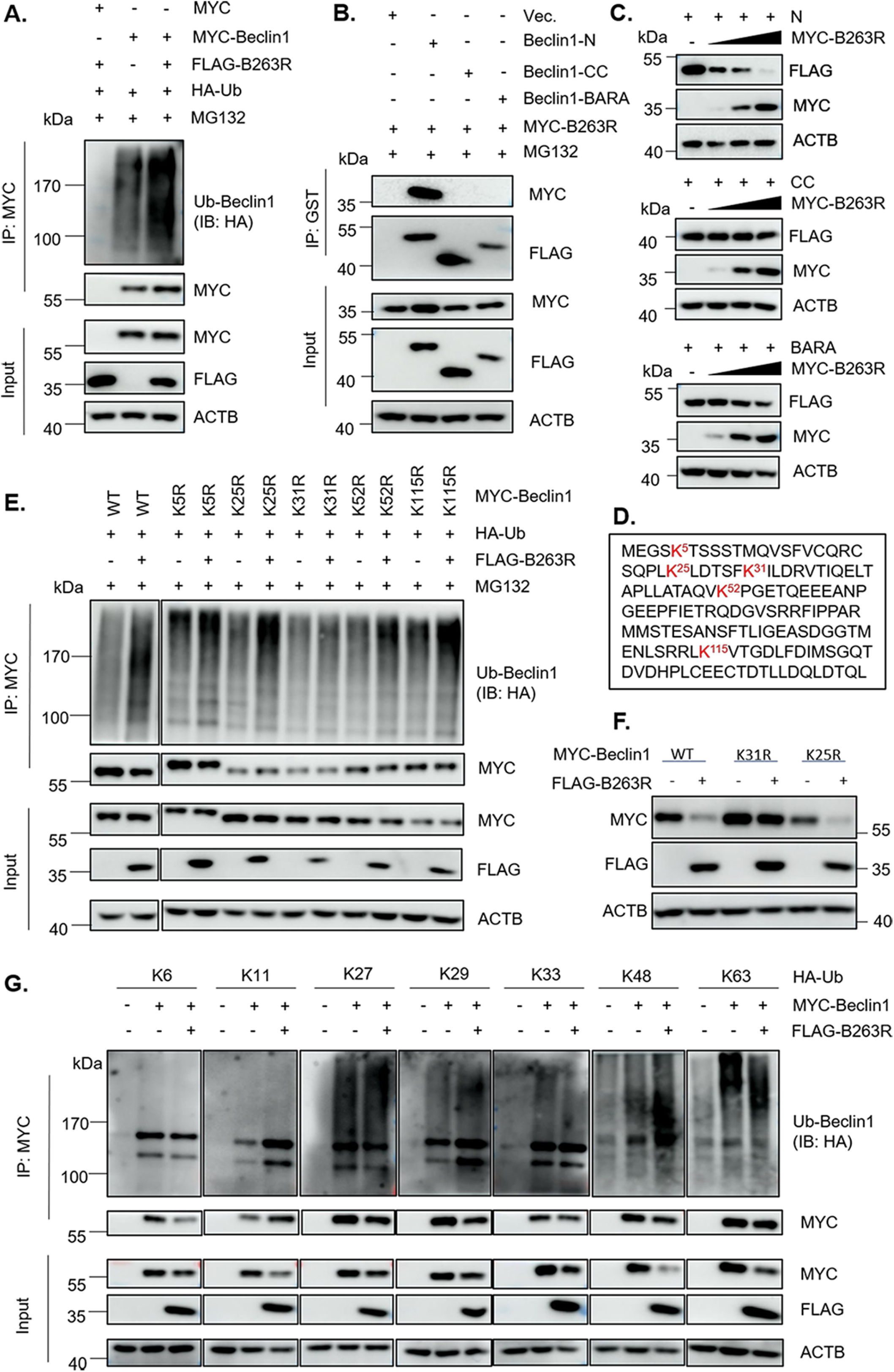
B263R promotes K48-ubiquitination of Beclin1 on K31. (A) B263R ubiquitinates the protein BECN1. MYC or the MYC-BECN1 vector was co-transfected with FLAG or the FLAG-B263R vector and HA-Ub plasmids into HEK293T cells for 36 h, followed by treatment with 20μM MG132 for 6 h. The cell lysates were then subjected to immunoprecipitation and immunoblot using the indicated antibodies. (B) MYC-B263R was co-transfected with FLAG-GST-tagged N-terminal region, CC domains, or BARA domain plasmids into HEK293T cells for 36 h, followed by treatment with 20μM MG132 for 6 h. The cell lysates were then subjected to immunoprecipitation and immunoblot using the indicated antibodies. (C) B263R promotes degradation of BECN1’s N-terminal region in a dose-dependent manner. Increasing amounts of B263R-expressing plasmids and FLAG-GST-tagged N-terminal region, CC domains, or BARA domain plasmids were co-transfected into HEK-293T cells, and the three domains were analyzed by detecting the protein levels. (D) The sequence of lysine residues of the N-terminal region on swine protein BECN1. (E) Vectors expressing MYC-BECN1 and its mutants, together with HA-Ub, FLAG or FLAG-B263R were co-transfected into HEK-293T cells for 36h, followed by treatment with 20μM MG132 for 6h. The lysates were subjected to immunoprecipitation and immunoblot with the indicated antibodies. (F) MYC-BECN1 or its two mutants were co-transfected with FLAG-B263R into HEK-293T cells for 36h, the immunoblot analysis was performed by blotting with anti-MYC, anti-FLAG, and anti-ACTB antibodies. (G) B263R mainly mediates K48-linked ubiquitination of BECN1. Vectors expressing MYC-BECN1 and FLAG-B263R together with HA-Ub mutants (K6, K11, K27, K29, K33, K48, and K63) were co-transfected separately into HEK-293T cells for 36 h. The lysates were immunoprecipitated with anti-MYC mAbs and immunoblotted using the indicated antibodies.

### Knockdown of BECN1 perturbs B263R mediated-autophagy

To better understand the involvement of BECN1 in B236R-mediated autophagy inhibition, we knocked down BECN1 by transfecting four individual siRNAs against BECN1 (siBeclin1#115, siBeclin1#664, siBeclin1#828, or siBeclin1#1222) or controll siRNAs (siNC) into the 3D4/21 cells for 48 h. As expected, the proteinand transcript level of BECN1 in the siBeclin1#1222 transfected cells were significantly lower than that in control siRNA (Figure 4A-B). The immunoblotting analysis of LC3 expression revealed that autophagy was inhibitive under the silence of swine BECN1 in Figure 4C. We then applied this knockdown of BECN1 on B263R expression cells. The results showed that the level of LC3-Ⅱ inhibition caused by B263R could be restored by silencing of BECN1 compared to negative control (Figure 4C). The data demonstrates that the BECN1 plays a crucial role in that B263R medicates autophagy inhibition.

**Figure 4.**
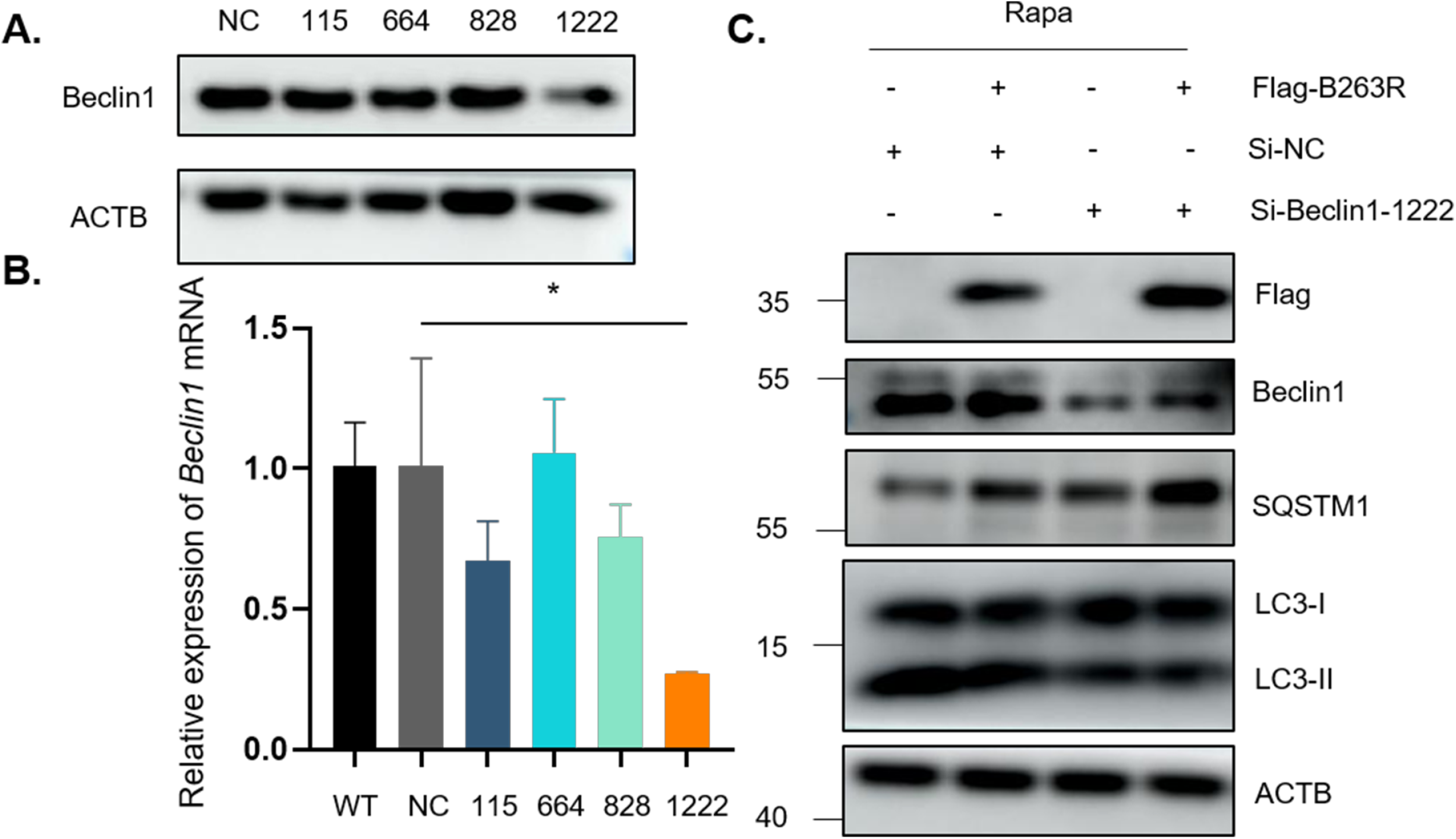
The siRNA knockdown of BECN1 affects B263R-mediated autophagy. (A) 3D4/21 cells were either mock transfected or transfected with 40 nM control siRNA or BECN1 siRNA for 48 h. Samples were analyzed by immunoblotting with anti-BECN1 and anti-ACTB antibodies. (B) Same samples in (A) were subjected to RT-PCR and qRT-PCR as described in the Materials and methods. (C) Under the siRNA knockdown of BECN1, 3D4/21 cells were transfected with FLAG or the FLAG-B263R, cells were treated with Rapamycin (5nM) for 4h. Samples were analyzed by immunobloting WB with specific antibodies as indicated.

### B263R inhibits the replication of adenovirus

Zhang et al. found that autophagy benefits for the replication of recombinant Adenovirus (rAdVs) (24). B263R Knockout failures to rescue the ASFV, we therefore validated the effect of B263R on virus replication by reconstructing rAdVs to express B263R. To address whether the rAdVs was impacted due to cellular toxicity, the cell viability was assessed by an WST-8 (2-(2-Methoxy-4-nitrophenyl)-3-(4-nitrophenyl)-5-(2,4-disulfophenyl)-2H-tetrazolium Sodium Salt) assay. The data showed that rAdVs expressing B263R was no toxic to the 3D4/21 cells at 1 MOI (Figure 5D). We then monitored the replication of rAdVs that contained B263R in 3D4/21 cells by observing the GFP fluorescent signals expressed by the virus. In comparison to the control rAdVs, the green signals of B263R decreased greatly (Figure 5A-B). The cells infected by rAdVs were subsequently harvested to analyze protein levels. As depicted in Figure 5C, GFP expression exhibited a significant decrease in comparison with those cells infected with the empty vector control rAdVs.

**Figure 5.**
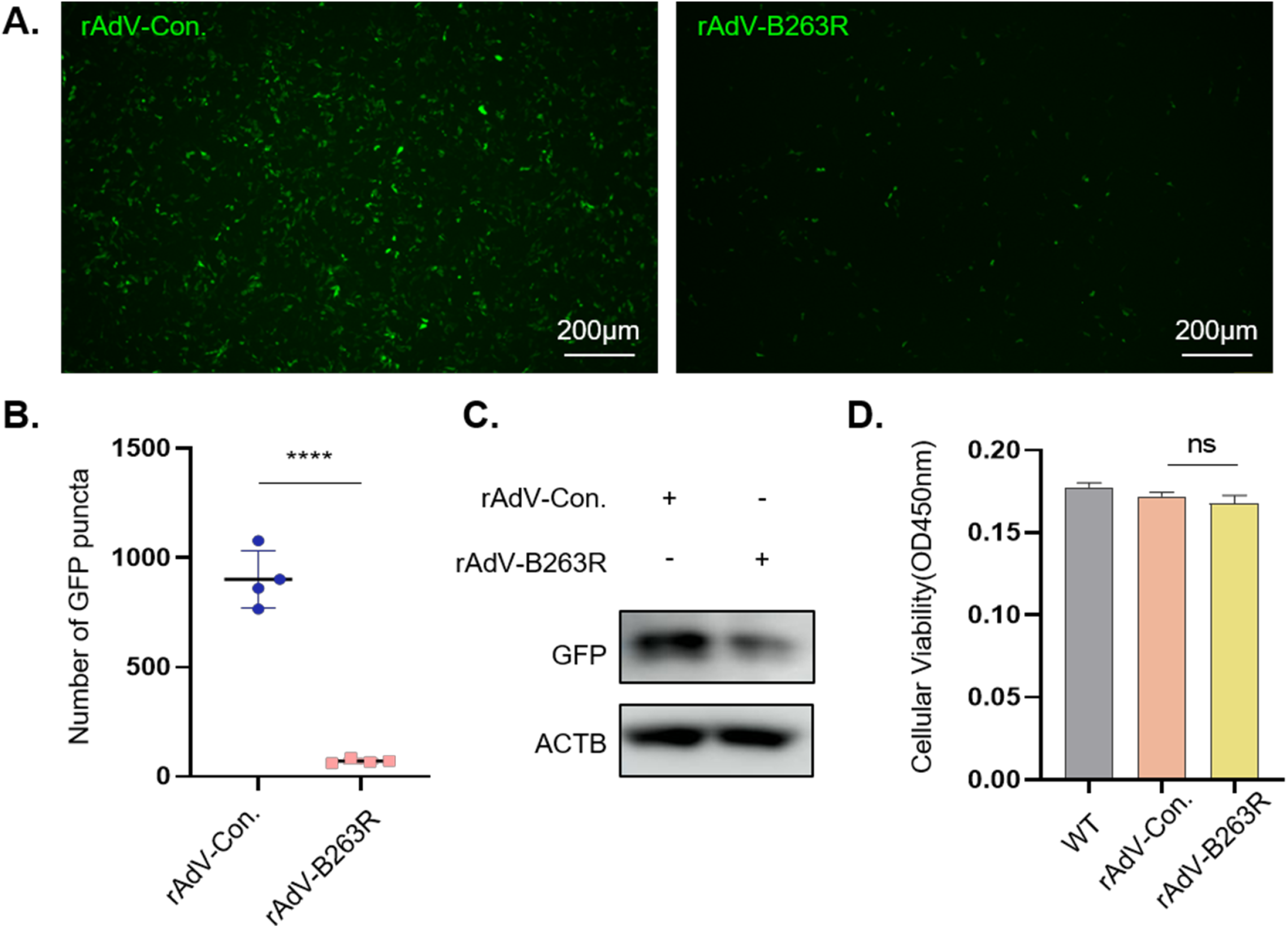
Effects of B263R on replication of adenovirus. (A) 3D4/21 cells were infected with GFP-tagged rAdVs and rAdVs expressing B263R at MOI of 1 for 24 h. The cells were then observed under a a fluorescence microscope. Scale bar: 200μm. (B) The numbers of puncta were counted by using Image J software. Error bars: Mean ± SD of 3 independent tests. two-tailed Student’s test; *p < 0.05; **p < 0.01; ***p < 0.001, ****p < 0.0001 compared to control. (C) The cells from A were harvested for immunoblot analysis by using anti-GFP, anti-ACTB antibodies. (D) Viability of cells infected with rAdVs and rAdVs expressing B263R at MOI of 1 for 24 h.

## Discussion

ASF is characterized by hemorrhage and increased mortality, causing significant economic setbacks to the global pig industry. There are currently no commercial vaccines against ASFV, which limiting the control of ASF worldwide. Recombinant ASFV with deletions of various virulence-related genes that induce a powerful effect, have received much attention in recent decades(25). Understanding the interplay between ASFV gene functions and host responses is crucial for elucidating virus characteristics and identifying potential targets for virus attenuation. Numerous studies have demonstrated that ASFV invasion triggers unfolded protein response (UPR) and apoptosis (26, 27). For instance, Zheng et al. conducted transcriptome profiling of swine macrophages infected with ASFV using single-cell RNA-sequencing technology (28). They observed downregulation of differentially expressed genes (DEGs) in ASFV-infected cells, particularly those associated with processes involving the autophagy mechanism and membrane localization. However, autophagy plays a crucial role in orchestrating the host cell’s response to stress, and it also represents a formidable threat selectively exploited by various viruses for inhibition (29). Previous studies have indicated that ASFV blocks the formation of autophagosomes induced by starvation (13). However, the expression of these autophagic genes in ASFV-infected pigs and the mechanisms by which they execute their functions have not been systematically studied.

BECN1, an autophagy gene in mammals, shares homology with the ATG6/Vps30 protein in yeast and plays a pivotal role in autophagy. It stimulates autophagy by modulating the activity of the lipid kinase Vps34 and facilitating the assembly of the BECN1-Vps34-Vps15 core complex, which is crucial for the localization of autophagic proteins to the pre-autophagosomal structure (30). The activity of BECN1 is tightly associated with the post-translational level such as phosphorylation, ubiquitination and acetylation (31-33). Ubiquitination, as a critical post-translational modification, participates in numerous cellular processes. Ubiquitin, a small protein with seven lysine sites, enables targeted proteins to undergo at least seven types of polyubiquitination. In our study, we observed that B263R interacts with BECN1 and mediates its K48-linked ubiquitination, resulting in BECN1 degradation via the proteasome. Notably, the analysis of BECN1 ubiquitination in Figure 3E indicates that the promotion of B263R on BECN1 ubiquitination is significantly reduced when the K^31^ residue is substituted with arginine. This finding aligns with the work of Tang et al.(34), who also found that K27-linked ubiquitination of BECN1 on K32 (exhibits homology with K31 in sus scrofa species) is relevant for BECN1 stability.

Adenoviruses are double stranded DNA viruses that exist in vertebrates (35). Due to their ability to mimic viral infections, AdVs are frequently employed in vaccine research and development (36, 37). They represent a valuable tool for assessing the impact of specific exogenous proteins on virus replication (24). AdVs induce autophagy by facilitating the conversion of LC3-I to LC3-II, with autophagy subsequently promoting viral replication (38). Our data reveal that the rAdVs expressing B263R effectively inhibits virus replication (Figure 5). This observation indicates a notable correlation between B263R’s inhibition of autophagy and its ability to suppress AdVs replication.

Taken together, these results suggest that B263R reduced autophagy activity by interacting with BECN1 and mediating K48-related ubiquitination of the K31 site. Moreover, the knockdown of BECN1 led to the restoration of LC3-II levels, indicating that B263R-induced autophagy inhibition involves BECN1 degradation. In our investigation, where adenovirus was employed to simulate DNA virus infection in cellular models, we noted a significant reduction in virus proliferation mediated by B263R. Further work needs to be evaluated whether the B263R-deleted ASFV could be a potential live attenuated vaccine in pigs. This research has provided deeper insight into the interaction between autophagy and ASFV, aiding in the development of targeted therapies and strategies to mitigate its impact on swine populations.

## Materials and methods

### Cell cultures and Adenoviruses

HEK-293T (CRL-11268) cells and PK-15cells (CCL-33) were cultured in Dulbecco’s modified Eagle medium (DMEM; Gibco, Carlsbad, CA USA) supplemented with 10% fetal bovine serum (FBS; 12B269, Uruguay). 3D4/21 cells were cultured in Roswell Park Memorial Institute (RPMI) 1640 (DMEM; Gibco, Carlsbad, CA USA) supplemented with 5% FBS. Adenoviruses expressing B263R (pAD100013-OE; Vigene Bio science, Jinan, China), and empty control were purchased from Vigene Biosciences Company (Jinan, China).

### Antibodies and reagents

Anti-ACTB (M1210-2) and anti-MYC (R1208-1) rabbit mAbs and anti-FLAG (0912–1) mouse polyclonal antibodies (pAbs) were obtained from Hangzhou HuaAn Biotechnology. Anti-LC3B (2775S) rabbit mAbs, and anti-ubiquitin (3936S) mouse mAbs were purchased from Cell Signaling Technology. Anti-SQSTM1 (18,420-1-AP) rabbit pAbs were provided by Proteintech. MG132 (HY-13259), CQ (HY-17589A) and Rapamycin (HY-10219) were purchased from MedChemExpress company. Earle’s Balanced Salt Solution (EBSS: 14155063) was purchased from Thermo Fisher Scientific.

### Western blotting

Cells were collected at specified time points and lysed immediately in lysis buffer containing 2% sodium dodecyl sulphate (SDS), 50 mM Tris-HCl, 1% Triton X-100, 150 mM NaCl with pH 7.5. The lysate proteins were separated though SDS-polyacrylamide gel electrophoresis (PAGE), and the proteins were then transferred onto nitrocellulose blotting membranes (NC) (10600001; GE Healthcare Life Science). The membranes were blocked with 5% non-fat dry milk and 0.1% Tween 20 for 30 min at 37°C, and then incubated with primary antibodies for 2h at 37°C, followed by incubating with horseradish peroxidase-conjugated anti-mouse/rabbit IgG (Kirkegaard & Perry Laboratories, Inc., 074–1506), finally imaged using AI680 Images (GE Healthcare, USA) after using enhanced chemiluminescence (ECL).

### DNA construction and transfection

B263R was amplified from gDNA of ASFV and cloned separately into vectors pCMV-Flag-N (635688; Clontech, Mountain View, CA, USA), pCMV-Myc-N (635689; Clontech, Mountain View, CA, USA), and pEGFP-C3 (#6082-1; Biosciences Clontech). BECN1 amplified from 3D4/21 cells was separately inserted into vector pCMV-Myc-N. The segments N-terminal, CC, BARA were amplified from BECN1 and cloned into pCMV-Flag-N-GST. Several mutants of MYC-BECN1 were constructed by site-specific mutation experiments. For cell transfection, HEK-293T, PK-15 or 3D4/21 cells were seeded on the designated plates according to the experimental scheme. When cells were 70-80% confluent, they were transfected with jetPRIME Transfection Reagent. After transfection with vectors for the specific time, the cells were harvested for subsequent detection.

### Immunofluorescence staining

The 3D4/21 cells were seeded on 35 mm glass-bottomed cell culture dishes for 24 h, and then transfected as indicated. After fixing using 4% paraformaldehyde for 10 min at room temperature, and permeabilized with 0.1% Triton-X-100 for 10 min. The cells were then blocked with 5% non-fat milk for 0.5 h, and incubated with the primary antibodies for 2 h at 37°C after washing three times with PBST, and with another three times of washing with PBST, the samples were then incubated with Alexa Fluor 546 (A10036, Invitrogen, USA) for 1 h at 37°C. After washing three times with PBST and stained with DAPI I (4’,6-diamidino-2-phenylindole) for 10 min to indicate nuclei. The cells were scanned with an LSM780 laser scanning confocal microscope (Zeiss, Oberkochen, Germany).

### Co-immunoprecipitation

The HEK293T cells were co-transfected with various plasmids for indicated time and then lysed using NP40 lysis buffer with phenylmethanesulfonylfluoride (PMSF, 1 mM) for 4 h at 4°C. Then, the supernatants from the cell lysis after centrifuging at 12,000 × g for 10 min, followed by incubation with anti-Flag antibody and Protein A/G beads for 4 h at 4°C. The supernatants were removed after centrifugation, and the pellets were suspended in washing buffer. This step was repeated for 3 times. The pellets were finally lysed in lysis buffer for WB analysis.

### Ubiquitin assays

The HEK293T cells were co-transfected with various plasmids for for 36h, followed by treatment with 25μM MG132 for 6 h and lysed with NP40 containing 6M urea and 1 mM PMSF for 4 h at 4°C. The lysates were then centrifuged at 12,000×g for 10 min at 4°C, the supernatants were diluted with NP40 and added to the Pierce Glutathione Agarose for 4 h at 4°C, the Pierce Glutathione Agarose was washed five times with NP40 containing 0.5 mM Dithiothreitol (DTT) (Sangon Biotechnology, A100281), and the supernatant was removed. Finally, the Pierce Glutathione Agarose was lysed with lysis buffer for western blotting.

### SiRNA knockdowns

SiRNAs targeting Beclin1 (siBECN1-115: 5′-GGAGCUUACAGCUCCAUUATT-3′, siBECN1-664: 5′-GGAGGAAGCUCAGUAUCAATT-3′, siBECN1-828: 5′-GUGGACAAUUUGGCACAAUTT-3′, siBECN1-1222: 5′-CGGCUCCUAUUCCAUCAAATT-3′ and control siRNA were synthesized by GenePharma (Shanghai, China). 3D4/21 cells were transfected with siRNAs using jetPRIME Transfection Reagent. Samples were harvested at the indicated time points for immunoblot analysis.

### Reverse transcription PCR (RT-PCR) and quantitative real time reverse transcription PCR (qRT-PCR)

Total RNA was extracted, treated with RNase-Free DNase, and reverse-transcribed into cDNA using a HiScript Ⅱ Q RT SuperMix (Vazyme Biotech Co., Ltd, R222). The resulting cDNA was then analyzed via qPCR using 2 × Taq Universal SYBG qPCR Mster Mix (Vazyme Biotech Co., Ltd, Q712). GAPDH expression served as an internal control for normalizing the relative abundance of Beclin1 mRNA. Beclin1-F: CTAAGACATCCAGCAGCAC, Beclin1-R: TTCAATAAATGGCTCCTCT. GAPDH-F: TGGTGAAGGTCGGAGTGAAC, GAPDH-R: GGAAGATGGTGATGGGATTTC.

### CCK-8 assay

The cell viability of 3D4/21 cells was measured by using an WST-8 assay (Beyotime, Cell Counting Kit-8, C0037). In brief, cells were seeded in 96-well plates at a density of 10^3^ cells per well in 100μl of medium. The plates were incubated in a 37°C incubator for 1h. The formazan dye was detectable by spectrophotometric analysis with OD450nm.

## ACKNOWLEDGMENTS

This study was supported by the National Natural Science Foundation of China (grant no. U21A20256) and and the Fundamental Research Funds for the Central Universities (2022-KYY-517101-0005).

## Figure Legends

**Figure S1. Analysis of ASFV-encoded proteins in autophagy**.

Screening of ASFV-encoded proteins for the regulation of autophagy in HEK293T cells by determining the protein levels of SQSTM1 and LC3-II.

**Figure S2. Analysis of host proteins that interacted with B263R**.

(A) HEK-293T cells were co-transfected with vectors expressing MYC or MYC-B263R and FLAG-Vps34 for 36h. At 36h post-transfection, cells were harvested for Co-IP. Immunoblot analysis was performed with anti-FLAG, anti-MYC, and anti-ACTB antibodies. (B) HEK-293T cells were co-transfected with vectors expressing MYC or MYC-B263R and FLAG-UVRAG for 36h. At 36h post-transfection, cells were harvested for Co-IP. Immunoblot analysis was performed with anti-FLAG, anti-MYC, and anti-ACTB antibodies. (C) HEK-293T cells were co-transfected with vectors expressing FLAG or FLAG-B263R and MYC-ATG14 for 36h. At 36h post-transfection, cells were harvested for Co-IP. WB analysis was performed by blotting with anti-FLAG, anti-MYC, and anti-ACTB antibodies.

